# T1R3 subunit of the sweet taste receptor is activated by D_2_O in transmembrane domain-dependent manner

**DOI:** 10.1101/2023.02.28.530404

**Authors:** Natalie Ben Abu, Yaron Ben Shoshan-Galeczki, Einav Malach, Masha Y. Niv

## Abstract

Deuterium oxide (D_2_O) is a water molecule in which both hydrogens are replaced by the heavier and rare isotope deuterium. We have previously shown that D_2_O has distinct sweet taste, which is mediated by the T1R2/T1R3 sweet taste receptor. Here we explore the effect of heavy water on T1R2 and T1R3 subunits. We show that D_2_O activates T1R3 transfected HEK293T cells similarly to T1R2/T1R3 transfected cells. The response to glucose dissolved in D_2_O is higher than to glucose dissolved in water. Mutations of Phenylalanine at position 730^5.40^ in the transmembrane domain of T1R3 to Alanine, Leucine or Tyrosine, impair or diminish activation by D_2_O, suggesting a critical role for T1R3 TMD domain in relaying the heavy water signal.

## 1. Introduction

Deuterium Oxide (D_2_O), commonly referred to as “heavy water’’ is a naturally occurring, but extremely rare molecule ^7^ that differs from regular water (H_2_O) by H-D isotopic substitution. This results in 10% increase in molecular weight, small differences in pH (+0.44 for D_2_O), and higher melting and boiling points ^5, 24, 27, 34^. Cellular function of mitosis was shown to be inhibited by D_2_O in animal and plant cells ^26^. In algae, D_2_O has been observed to affect membrane depolarization and activation of Ca^2+^ channels ^1^. Protozoa, bacteria and yeasts are able to tolerate 70% -100% D_2_O ^22^. In whole organism studies, small amounts of D_2_O, usually lower than 20% of body weight, were tolerated well over short time periods ^36^. In fact, D_2_O and deuterated drugs are used in drugs metabolism studies in humans and other animals, and recently also as novel versions of existing drugs ^33^. Effects of deuteration on drugs in enzymatic reactions ^38^, as well as on interactions with receptors, are studied using cell-based and computational approaches ^16, 21^. For example, a recent molecular dynamics simulations study indicates that D_2_O has a higher ability to form water−water hydrogen bonds than water−amino acid hydrogen bonds ^32^. However, detailed effects of D_2_O on protein function remain to be explored.

Here we investigate the effect of heavy water on the human sweet taste receptor, T1R2/T1R3, in cell-based functional assays. The debate regarding the potential taste of heavy water has begun almost 90 years ago ^15, 34^. We have recently shown, using an integrated sensory, cell-based and computational approaches study, that humans differentiate between D_2_O and H_2_O based on taste alone, that D_2_O is sweeter than same-purity H_2_O and that D_2_O may add to the perceived sweetness of sweeteners ^4^. Recognition of sweetness is related to T1R subfamily which contains three taste receptors; T1R1, T1R2, and T1R3. The main sweet taste receptor is the T1R2/T1R3 heterodimer of Class C GPCRs ^23, 40^, though stimulation of sodium-dependent glucose transporter (SGLT1) expressed in sweet taste may also participate in signal transduction of sugars ^25^. Lactisole, a known sweetness inhibitor acting via the T1R3 monomer of the T1R2/T1R3 ^18^, suppressed the sweetness of D_2_O for humans, suggesting T1R2/T1R3 as D_2_O target. Accordingly, T1R2/T1R3-transfected HEK293T cells were activated by D_2_O and this cellular response was also inhibited by lactisole ^4^. Interestingly, similarly to the lack of murine response to the artificial sweetener cyclamate ^2^, mice did not show preference for D_2_O over regular water, raising the hypothesis that the site of interaction of D_2_O with the receptor may be close to the binding site of cyclamate ^4^.

T1Rs proteins have three main domains: the large N-terminal “Venus flytrap” domain (VFT) which was shown as the orthosteric site for sugar binding by numerous site-directed mutagenesis studies ^13^; the Cysteine rich domain (CRD) that contains nine highly conserved cysteines and is targeted mainly by sweet proteins, and the seven helical transmembrane domain (TMD)^6^. The TMD binding site is analogous to the orthosteric site in Family A GPCRs, and in human hT1R3 was shown to accommodate cyclamate, with F730 (5.40 BW position ^3^ of the T1R3 (http://www.gpcrdb.org/)) established as important for activation ^19^. Previous highlighted the impact of T1R2 and T1R3 single nucleotide polymorphisms (SNPs) on consumption of sugars as well as on sweet products ^11, 29^. Examining activation of the receptors expressed in HEK293T cells, Dubovski et al. showed differences in sensitivity comparing two sequences of T1R2: the older reference sequence (NM_152232.1) which contains four of the SNPs - I191V, R317G, I486V and S9C and an updated reference sequence (NM_152232.4). Residue R317 in the VFT domain of T1R2, was shown to be responsible for this difference. Furthermore, they showed that site-directed mutagenesis of Serine at position 147 in the binding site of T1R3 VFT domain to Alanine, abolished receptor activation by D and L glucose enantiomers ^10^.

Interestingly, T1R3 without T1R2 (possibly functioning as a homodimer) was shown to respond to monosaccharides and disaccharides ^10, 39^. This is of physiological relevance, since T1R3 (but not T1R2) expression was shown in the gastrointestinal tract ^25^ pancreas and liver ^9, 31^.

Here we investigate whether T1R2 or T1R3 may react to D_2_O when expressed without the other unit, and test our hypothesis regarding the key role of T1R3 TMD domain in sensitivity to heavy water.

## 2. Experimental Procedures

### Materials

Cyclamate, D-glucose, powder Dulbecco’s Modified Eagle’s Medium (DMEM), were purchased from Sigma-Aldrich (St. Louis, Missouri, United States). Deuterium Oxide (D_2_O) was purchased from Tzamal D-Chem Laboratories Ltd (Kiryat-Matalon, Petach Tikva, Israel). Unless otherwise noted, the typical concentration of tested solutions was as follows: cyclamate (0.01M), D_2_O (49.9M) and D-glucose (0.96M).

### Cell Culture and IP - One HTRF Assay

Human embryonic kidney 293T (HEK293T from ATCC) cells were cultured, maintained, and transiently transfected as described previously ^4^. All treatments were done in triplicate, and all experiments were repeated at least three times. In general, cells were grown to a confluency of approximately 85-90 % and transiently transfected with plasmids encoding T1R proteins (T1R2, T1R3) and Gα16gust44 (a chimeric G protein and subunit containing the last 44 amino acids of gustducin). After 24h, cells were seeded in 24-well culture plates (0.5 ml cells per well) and maintained for 8-12 h at 37 °. Next, cells were “starved” overnight by changing the medium to 0.1% DMEM (containing 0.1% FBS) without glucose, aiming to reduce the basal activity of the cells. After an additional 18 h, cells were exposure to tested compounds by addition of 0.5ml of each with 50 mM Lithium Chloride (LiCl) needed for IP-One accumulation (based on Cisbio IP-One HTRF assay) and dissolved in 0.1 % DMEM (without glucose), for 5 minutes. In case of monitoring the effect of heavy water, 0.1 % DMEM medium was made using D_2_O instead of D_2_O water. Upon exposure time, tastant solution was replaced with fresh medium (0.1 % DMEM) containing 50 mM LiCl for another 1h and then washed with 100μl cold phosphate buffered saline (PBS) + Triton X-100, and kept at -80◦C for at least 30min, in order to dissolve the cell membrane. Cell lysate was mixed with the IP-One HTRF detection reagents (IP1-d2 conjugate and Anti-IP1 cryptate TB conjugate), and added to each well in a 384-well plate for 60min incubation at room temperature. Changes in IP-One levels were read using Clariostar plate reader (665nm/620nm emission ratio).

### Statistical analysis

Dose – response curves were fitted by non-linear regression using the algorithms of PRISM 7 (GraphPad Software, San Diego, CA, USA). All responses are presented as the means ± SEM of IP1 accumulation (%) and were normalized based on maximum response of the full sweet taste heterodimer receptor T1R2/T1R3 to 0.96M glucose (Emax=100%), as well as tested in comparison to basal activity.

### Mutants

Site-directed mutagenesis in T1R3 was performed by Transfer-PCR (TPCR), as described previously ^12^. All mutants were activated by glucose, indicating that amino acids substation at these positions did not affect protein expression.

### Water mapping

Mapping of water molecules was calculated on the AlphaFold2 ^20^ model of hT1R3 (Uniprot ID: Q7RTX0). The model was downloaded from the AlphaFold2 EBI database (https://alphafold.ebi.ac.uk), and then prepared and minimized with Schrödinger Maestro protein preparation panel (Maestro Version 12.7.161, MMshare Version 5.3.161, Release 2021-1, Platform Windows-x64). The minimized receptor model was then used to map possible water molecule positions near F730 using OpenEye SZMAP with default settings (SZMAP 1.6.5.2: OpenEye Scientific Software, Santa Fe, NM. http://www.eyesopen.com, OpenEye Applications & Toolkits 2022.1.2).

## 3. Results

### D_2_O activates the T1R3 subunit

To assess which of the T1R subunits of the heterodimeric human sweet receptor is required for sensing D_2_O. Hence, we expressed separately each one of the human T1R subunits; T1R2 or T1R3, by transient transfection in HEK293T cells, along with Gα16gust44. Activation of the receptor was monitored by IP1 accumulation. 960mM of D-glucose was used here as control for each transfection, based on previous work _4, 10_.

As seen in Figure 1A, T1R3 expressed without T1R2 responded to D_2_O similarly to the response of T1R2/T1R3 whereas the T1R2 expressed without T1R3 did not (p.v ≤0.0001). As expected, D-glucose caused IP1 elevation in all types of transfections. Figure 1B illustrated a similar experiment in functional cell assay: we find that 960mM D-glucose dissolved in D_2_O elicited significantly higher (p.v ≤ 0.005) IP1 values compared to the same concentration dissolved in H_2_O, whereas no significant difference was observed for cyclamate. This result raises the possibility that cyclamate and heavy water share a common binding region.

**Fig. 1.**
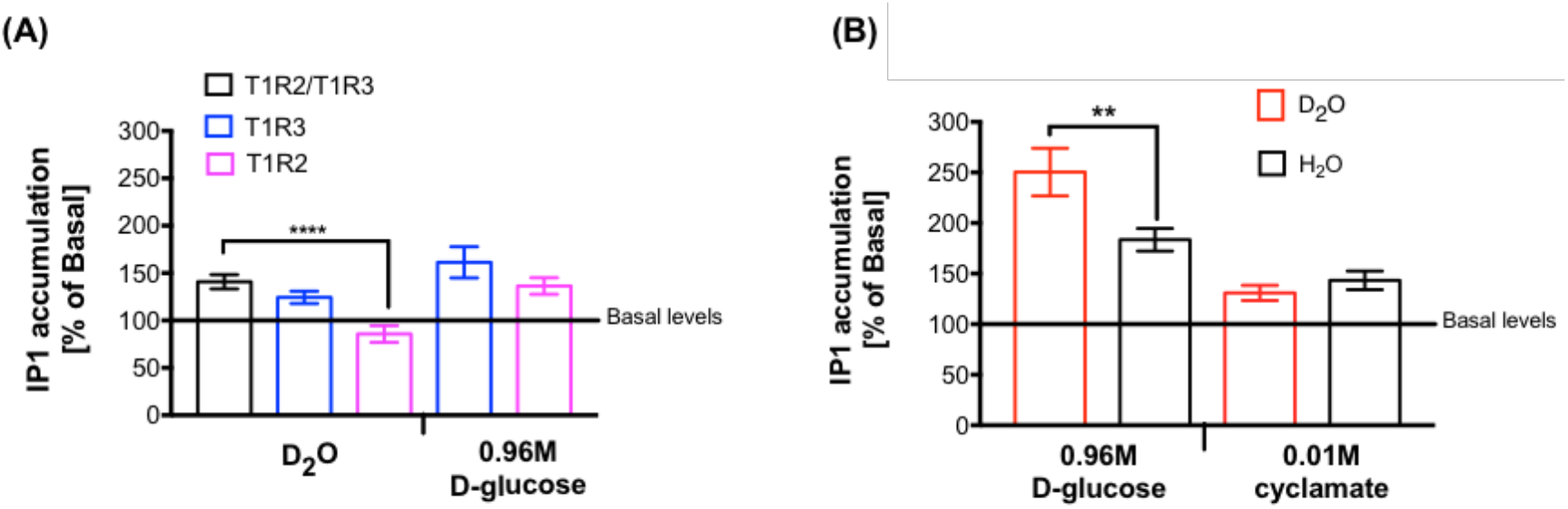
D_2_O signaling requires T1R3 subunit. **(A)** The receptors composed of T1R2(NM_152232.1)/T1R3(black), T1R3(blue) or T1R2(NM_152232.1, pink) were co-expressed by transient transfection in HEK293T cells along with Gα16gust44 and tested toward their responses to D_2_O in comparison to the positive control of D-glucose. **(B)** The effect of D_2_O (red) on T1R2/T1R3 activation elicited by different sweet molecules. On y-axis, changes in IP1 accumulation upon stimulus application are shown as % of basal – pure H_2_O; x-axis,100% D_2_O, 0.96M glucose or 0.01M cyclamate. Asterisks indicate IP1 changes that are significantly different from basal (* for p ≤ 0.05, ** for p ≤ 0.005 and **** for p ≤0.0001) using t-test. All tested solutions were done in at least triplicate; each experiment (transfection) was repeated three to five times.

We next tested varying concentrations of heavy water for different types of transfections. D_2_O dose response curves were obtained for cells expressing T1R3 (Emax=50% of activation by 0.96M glucose, which was used as reference), while cells transfected with T1R2 subunit were not activated (Fig. 2A). Dubovski et al. showed that the T1R2 (NM_152232.4) is more sensitive to presence of D-glucose ^10^. In order to understand whether these SNPs affect the sensitivity of activation by D_2_O, we compared the dose response curves of HEK293T cells, transfected with either the T1R2 (NM_152232.1) or the more sensitive version (NM_152232.4), along with T1R3. As shown in Fig. 2B, we observed only minor differences in responses to D_2_O by these two versions of T1R2 (NM_152232.1 Emax = 60% and Emax = 50% for NM_152232.4), confirming that T1R3 subunit play the dominant role in D_2_O activation.

**Fig. 2.**
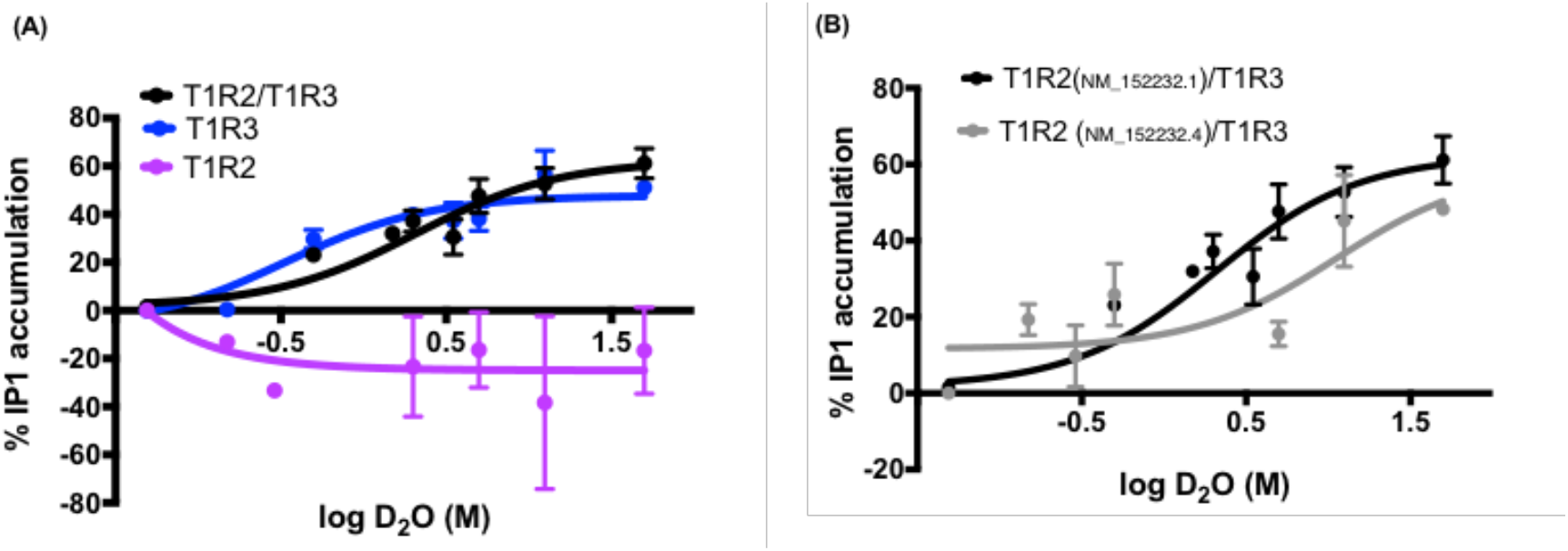
D_2_O dose response curve characterization. **(A)** T1R3 transfected HEK293T cells achieved D_2_O dose-responsive manner. Varying concentrations of deuterium Oxide were tested among different transfections; T1R2(NM_152232.1)/T1R3(black), T1R3/T1R3(blue) or T1R2/T1R2(NM_152232.1, pink). **(B)** Dose-response curves of HEK293T cells transiently transfected with T1R2(NM_152232.1)/T1R3 (black); or T1R2(NM_152232.4)/T1R3 (grey); All data were normalized based on maximum response of the full sweet taste heterodimer receptor ^10^T1R2/T1R3 to 0.96M glucose. All tested solutions were done in at least triplicate; each experiment (transfection) was repeated three to five times.

### Transmembrane domain of hT1R3 is involved in D_2_O response

Once we have established that the T1R3 is essential and sufficient for activation by D_2_O, we aimed to identify the responsible sub-domain in T1R3. First, we tested the effect of alanine mutations in residues: 730 (F730A, the TMD domain) and 147 (S147A, VFT domain, see Fig 3A for location on the receptor and 3B for human-mouse sequence alignment in the relevant regions on T1R3 activation by D_2_O. T1R3 plasmids of T1R3F730A or T1R3S147A were transfected together with Gα16gust44 (without T1R2), and compared to WT T1R3. (Fig. 3C). None of the mutants was activated by D_2_O; yet IP1 levels for F730A were significantly lower than WT basal activity (p.v≤ 0.0001) while the activation of S147A was similar to WT basal activity. Importantly, cells transfected with either one of the alanine mutants of T1R3, were activated by 0.96M of D-glucose (p.v<0.05; Fig. SI.2).

**Fig. 3.**
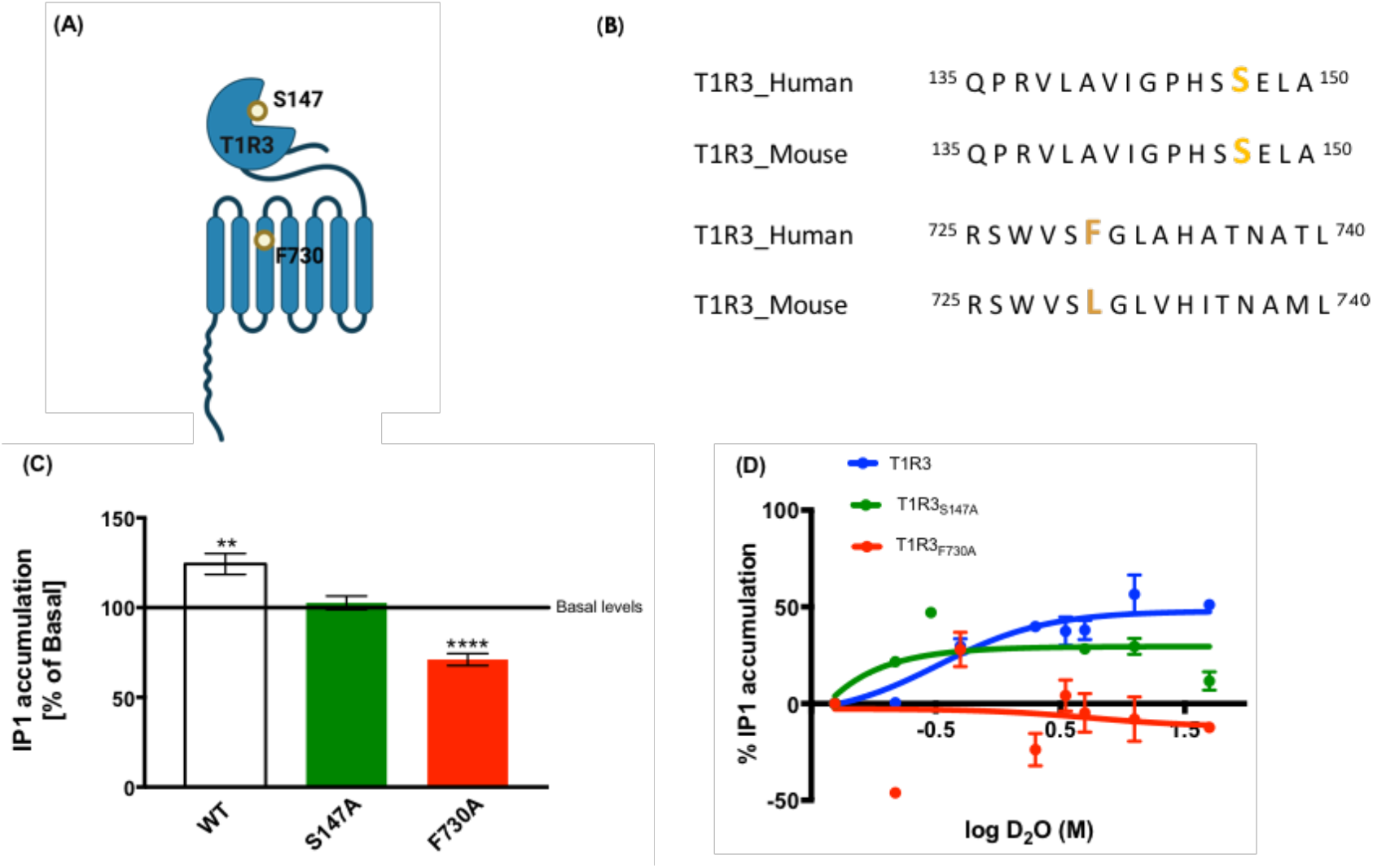
Involvement of TM helix 5 of hT1R3 in D_2_O binding. **(A)** Schematic picture of T1R3 subunit with domain-specific tested residues. **(B)** The alignment of human and mouse T1R3 sequences with an emphasis (orange) on the tested residues. **(C)** The effect of T1R3 F730A or T1R3 S147A mutant on activation by 100% D_2_O. T1R3 mutated human-receptors were co-expressed by transient transfection in HEK293T cells along with Gα16gust44. Asterisks indicate IP1 changes that are significantly different from basal levels (\**p* ≤ 0.05 \*\**p* ≤ 0.005 and \*\*\*\**p* ≤ 0.0001) using *t*-test. **(D)** The effect of alanine mutants in putative D_2_O-binding areas in the hT1R3 on D_2_O response. All data were normalized based on maximum response of the full sweet taste heterodimer receptor T1R2/T1R3 to 0.96M glucose.

As seen in dose response curves in Fig. 3D, both F730 and S147A mutants dramatically reduced Emax values, but cells transfected with F730A mutant had completely abolished the activation by D_2_O, while S147A mutant was still active.

### Exploring the 5.40 position

S147A mutant retained significant responsiveness to D_2_O, while F730A mutant showed no activity. The alignment of human and mouse T1R3 sequences shows that, at position 147 (VFT domain) both human and mouse have Serine, while at position 730 (5.40 TMD differs between these two species (Fig. 3B). We have also established a different behavioral effect to D_2_O of human and mice ^4^ and therefore focused on residue 730. In human, this residue is an aromatic amino acid Phenylalanine, which is predicted to interact with additional aromatic residues (Fig. 4A), while in mice this position harbors Leucine. Hence, we aimed to check the importance of aromaticity in this area. We mutated Phenylalanine (aromatic and hydrophobic) to Leucine (aliphatic and hydrophobic) and Tyrosine (aromatic). F730L mutant showed a complete loss of responsiveness to D_2_O, similarly to F730A, while F730Y still had some activity (Emax=20%) (Fig. 3B). We next confirmed that the effect of 730 T1R3 residue remains also in the framework of the full transfection (T1R2/T1R3) as seen in Figure 3C.

**Fig. 3.**
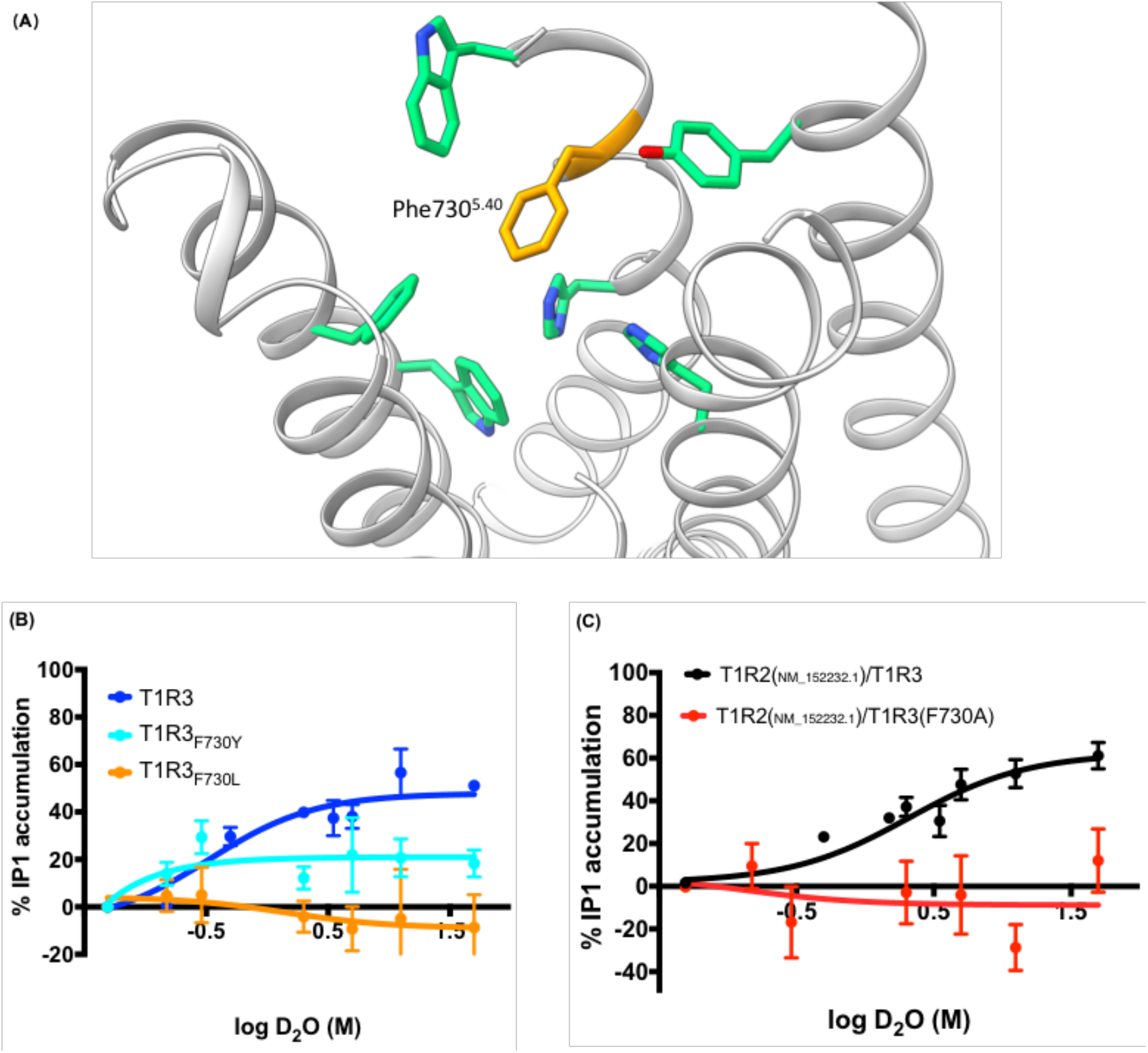
The importance of residue 730 in TMD pocket of T1R3. **A**. Snapshot of potential biding area of D_2_O to T1R3 based on AlphaFold model in which we found out that surrounding residues (within 4 angstrom) of F730^5.40^ are mainly aromatic amino acids. **B**. Dose-response profiles toward D_2_O of T1R3 F730^5.40^ mutants – substitution to tyrosine (cyan) and leucine (orange). **C**. The effect of F730A mutant of T1R3 on full transfected cells; T1R2(NM_152232.1)/T1R3(black), T1R2/T1R3F730A (red). T1R3 mutated human-receptors were co-expressed by transient transfection in HEK293T cells along with Gα16gust44 alone or with T1R2 (NM_152232.1) for full transfected cells. All data were normalized based on maximum response of the full sweet taste heterodimer receptor T1R2/T1R3 to 960mM glucose

## 4. Discussion

In this study, we continued to characterize the effects of D_2_O on human sweet taste receptor. This followed our previous work that showed D_2_O is mildly sweet to humans (and not to mouse) and that this sweetness is mediated via T1R2/T1R3 heterodimer ^4^. In the present study, we showed that D_2_O can also active T1R3 expressed HEK293T cells without the T1R2 monomer. The responses of HEK293T cells were characterized and showed similar dose-response curves and Emax values. The more sensitive variant of T1R2 (NM_152232.4) did not affect the D_2_O-elicited dose response. These outcomes pointed to T1R3 as the essential subunit for receptor activation by D_2_O. Furthermore, D_2_O had an added effect to the response elicited by 0.96M D-glucose, which had been shown as recognized by the VFT domain of T1R3 ^10^. However, D_2_O did not have the same effect on the activation elicited by 0.01M cyclamate, a sweetener that had been shown to be detected by the TMD of T1R3 (Jiang, Cui et al. 2005). This is similar to our previous finding that mice were not attracted to heavy water ^4^. Together, these indications supported the hypothesis that the interaction site of D_2_O with the T1R3 receptor should be near the binding site of cyclamate in the TMD of T1R3.

Here, we measured D-glucose to define which mutations (F730A in the T1R3 TMD or S147A in the T1R3 VFT) may cause lower activity. Both mutants were activated by D-glucose, thus, indicating that T1R3 remained an active protein, but to a lower degree, even though D-glucose is recognized by VFT domain, but probably not the TMD of T1R3. The indirect effect could be explained by reduced trafficking to the cell membrane, reduced receptor stability ^14, 30, 37^ or indirect allosteric effects ^28^. Importantly, our experimental results showed that for D_2_O, mutation F730A completely abolished activation by D_2_O, while S147A remained active, albeit with dramatically reduced Emax value. We have therefore concluded that the TMD region of T1R3 is critically important for the activation of this receptor by heavy water.

A CryoEM structure of class C orphan GPCR, GPR15, was recently solved ^17^. Residue D579^5.40^, equivalent to F730^5.40^ in T1R3, as well as in other class C GPCRs, belongs to ionic locks network and takes part in stabilization of transmembrane region. This position had been shown to be part of the allosteric binding sites in class C GPCR metabotropic glutamate receptor 5 ^8, 28^. Due to different residues in position 730 of T1R3 in human (Phenylalanine - aromatic and hydrophobic) and in mouse (Leucine - non-aromatic and hydrophobic), we explored additional mutations there. Indeed, F730L mutation completely abolished the response to D_2_O, just like the F730A mutation. F730Y however, had a similar effect to S147A, with only reduced D_2_O activity. F730A and F730L mutations also abolished the response to cyclamate ^19^.

Computational water mapping near residue F730, indicated that water molecules can potentially get into the TMD pocket (Fig. SI2). This is in accord with other studies of GPCRs that confirmed that many of these receptors have transmembrane water molecules ^35^. Hence, substitution of normal water molecules with D_2_O in this region could lead to stabilization of the active conformation, thus explaining the difference in receptor activity compared to regular water. Furthermore, both proteins and biomembranes tend to be slightly more compact and rigid in D_2_O that has stronger hydrogen bonding, than in H_2_O ^32^, suggesting a possibility for additional indirect effects of heavy water. How exactly heavy water affects T1R3 conformations, and how the effects differ in other T1Rs, present intriguing topics for further studies.

In summary, here we demonstrate that heavy water elicits functional response of the T1R3 subunit, but not through the T1R2 subunit of the sweet taste receptor, and that T1R3 activation depends on the residue found in position 5.40. Our findings are important since heavy water is used in clinical procedures, and T1R3 is expressed not only on the tongue but also in extra-oral tissues. Moreover, elucidation of effects of D_2_O on some but not other Family C GPCRs will help in understanding of the role of water in family C GPCRs function.

## 5. Acknowledgments

The authors thank Dr. R. F. Margolskee for the pcDNA3 of chimeric Gα16gust44and Dr. Maik Behrens for the pcDNA3 of T1R3 and for the pcDNA5FRT PM of T1R2S9C, I191V, R317G, I486V. We thank Dr. Yoav Peleg for help with sequencing and site directed mutagenesis; and for the construction of pcDNA5 of NM_152232.4, pcDNA5 of T1R3F730A, pcDNA5 of T1R3 F730L, pcDNA5 of T1R3F730Y, and pcDNA3 of TIR3S147A, and Prof. Michael Naim for helpful discussions. The study was supported by ISF grant #1129/19.

