## Supplemental figures for "T1R3 subunit of the sweet taste receptor is activated by D_2_O in transmembrane domain-dependent manner"

### Supplementary information

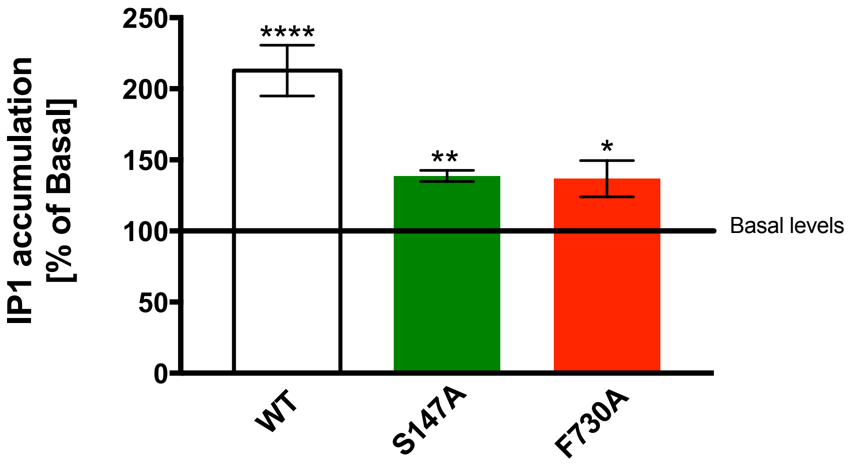


**Fig. SI1.** The effect of T1R3 F730A or T1R3 S147A mutant on activation by 960Mm D-glucose T1R3 mutated human-receptors were co-expressed by transient transfection in HEK293T cells along with Gα16gust44. Asterisks indicate IP1 changes that are significantly different from basal levels (*$p \leq 0.05$ **$p \leq 0.005$ and ****$p \leq 0.0001$) using *t*-test.

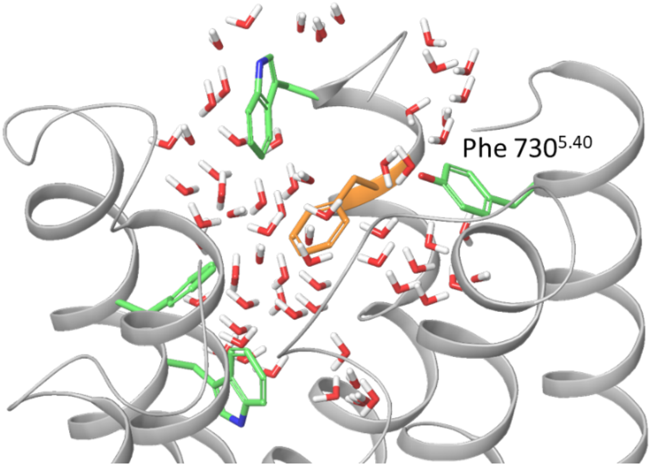


**Fig. SI2. Predicted water molecules near residue 730 in TMD pocket of T1R3. A.** Snapshot of predicted water molecules in the potential biding area of $D_2O$ to T1R3 based on AlphaFold model and surrounding aromatic amino residues (within 4 angstrom) of F730$^{5.40}$.
